# MiRformer: a dual-transformer-encoder framework for predicting microRNA-mRNA interactions from paired sequences

**DOI:** 10.1101/2025.11.21.689769

**Authors:** Jiayao Gu, Can Chen, Yue Li

**Affiliations:** School of Computer Science, McGill University, Quebéc, Canada; Mila - Quebéc AI Institute, Quebéc, Canada

**Keywords:** microRNA, Transformer, Post-transcriptional regulation

## Abstract

**Motivation:** MicroRNAs (miRNAs) regulate gene expression by binding to target messenger RNAs (mRNAs), inducing translational repression or mRNA degradation. Accurate prediction of miRNA–mRNA interactions and precise localization of binding and cleavage sites are critical for understanding post-transcriptional regulation and for enabling RNA therapeutics. Existing computational methods often rely on handcrafted or indirect features, scale poorly to kilobase-long mRNA sequences, or provide limited interpretability.

**Results:** We present `MiRformer`, a transformer-based framework that jointly predicts miRNA–mRNA interactions and localizes miRNA binding and cleavage sites directly from raw sequence pairs. MiRformer employs a dual-transformer encoder architecture for miRNA and mRNA sequences and incorporates a sliding-window attention mechanism to efficiently model kilobase-long mRNA contexts while preserving nucleotide-level resolution. Across multiple benchmrks, MiRformer achieves state-of-the-art performance on interaction prediction, binding-site localization, and cleavage-site identification from experimental Human Degradome-seq data. Beyond accuracy, MiRformer provides strong interpretability: attention patterns consistently highlight miRNA seed regions within 500-nt mRNA windows, revealing clear and biologically meaningful interaction signals. When applied to jointly infer binding and cleavage sites across 13k miRNA–mRNA pairs, predicted sites frequently co-localize, supporting a miRNA-mediated degradation mechanism.

**Availability:** Python code and datasets are publicly available at https://github.com/li-lab-mcgill/miRformer.

**Contact:** Please contact corresponding author at

## 1. Introduction

MicroRNAs (miRNAs) are small endogenous non-coding RNAs of approximately 22 nucleotides that regulate gene expression by binding to target messenger RNAs (mRNAs), leading to their degradation or translational repression [Bartel, 2004]. Each miRNA can target hundreds of mRNAs, forming complex post-transcriptional regulatory networks that control diverse biological processes such as cell proliferation, differentiation, and apoptosis. Dysregulation of miRNA expression has been implicated in numerous diseases, including cancer and neurological disorders. Accurate prediction of miRNA–mRNA interactions is therefore crucial for understanding gene regulatory mechanisms and for advancing RNA-based therapeutics. However, identifying true target sites remains highly challenging, as many functional interactions involve non-canonical seed regions that deviate from perfect Watson–Crick-Franklin base pairing [Hejret et al., 2023], complicating the interpretation of miRNA targeting rules and motivating the development of more accurate computational models.

Although experimental assays and next-generation sequencing approaches can map miRNA–mRNA interactions, they remain labor- and time-intensive, motivating the development of accurate in-silico predictors. Traditional heuristic tools such as TargetScan scan the 3^′^UTRs of target mRNAs for conserved seed matches across species [Agarwal et al., 2015]. While effective for canonical sites, these approaches rely on predefined alignment and conservation features, making them difficult to generalize to novel binding patterns or scale to kilobase-long sequences. To improve accuracy, deep learning methods such as DeepMirTar and miRAW incorporate handcrafted biological features—including minimum free energy, base pairing probability and sequence stability —alongside raw sequences [Wen et al., 2018, Pla et al., 2018]. However, these features are either derived from experiment data or are sensitive to small perturbations to input sequences, such as single-nucleotide variation. When used as precomputed input to model, such features may introduce variability and complicate end-to-end learning. The miTAR framework is introduced as an advanced system utilizing a sophisticated hybrid deep learning model [Gu et al., 2021]. This model integrates convolutional neural networks with recurrent neural networks to significantly enhance the capacity for recognizing patterns within biological data. However, this capability is specifically limited to analyzing short concatenated sequences of miRNA and mRNA, which presents a constraint in its application scope. REPRESS consists of a stack of residual connections between convolutional layers that infers miRNA binding and Degradome-seq read counts from the mRNA sequence [Kanuparthi et al., 2025]. However, when applied to human and mouse data across 39 cell lines and tissues, the model does not take the miRNA sequence as input and instead predicts only miRNA target sites that are highly expressed in the specific cell lines and tissues under study, which limits its ability to generalize to novel miRNA target sites. Overall, existing methods mainly suffer from two fundamental limitations: reliance on manually crafted features derived from limited experimental data and poor scalability to kilobase-long mRNA sequences.

Recent advances in transformer models have revolutionized biological sequence modeling by capturing both global and local contextual features through attention mechanisms [Avsec et al., 2021, Wang et al., 2024, Dalla-Torre et al., 2024, Patel et al., 2025]. In particular, transformer-based RNA language models are highly effective at capturing miRNA-mRNA interactions from raw sequences. Mimosa leverages two independent encoders - one encodes mRNA sequence and the other encodes miRNA sequence, and discovers local target sites by Smith-Waterman local alignment. This approach is a unique combination of both advanced deep-learning and classic alignment algorithm [Bi et al., 2024]. RNAErnie, a transformer-based RNA language model pretrained on multi-level motif-aware masking strategies, outperforms both heuristic methods and other transformer-based RNA language models on predicting miRNA-mRNA interactions [Wang et al., 2024].

In this study, we present `MiRformer`, a hybrid convolution and transformer framework that addresses three key limitations of existing miRNA target prediction methods: (1) inability to scale beyond short, fixed-length mRNA inputs—MiRformer employs sliding-window self-attention [Beltagy et al., 2020] in a dual-encoder architecture, scaling linearly to full-length 3’UTRs; (2) lack of a unified model for joint prediction of target, seed locations and cleavage sites—MiRformer simultaneously predicts three tasks with high accuracy, while convolutional tokenization with seed-length kernels produces interpretable attention patterns that consistently highlight seed regions; and (3) signal dilution in long sequences—MiRformer introduces a novel sliding-window cross-attention with Log-Sum-Exponential (LSE) pooling amplifies sparse seed signals during window aggregation, maintaining precise localization even when the seed occupies less than 1% of the input.

Through comprehensive benchmarking, MiRformer achieves state-of-the-art performance on all tasks, consistently outperforming existing methods. Despite never observing jointly annotated seed and cleavage sites during training, it produces biologically plausible co-localized predictions when trained separately on TargetScan seed labels and Degradome-seq cleavage data. To validate generalizability, we evaluate on miRBench [Sammut et al., 2025], an independent benchmark derived from experimental CLASH and chimeric eCLIP data, where MiRformer achieves competitive performance without retraining. To our knowledge, MiRformer is the first framework to jointly predict miRNA–mRNA interactions, seed binding sites, and cleavage sites at nucleotide resolution while scaling efficiently to full-length mRNA sequences.

## 2. Method and Materials

### 2.1 MiRformer model details

#### Convolution layers to extract initial sequence features

All miRNA and mRNA sequences are tokenized at the nucleotide level. To enhance local continuity, the ribonucleotide tokens are first processed through two convolution layers with kernel sizes of 5 and 7, respectively (Fig. 1a). This design is inspired by RNAGenesis [Zhang et al., 2024], which demonstrated that convolution-augmented embeddings improve token coherence.

**Fig. 1.**
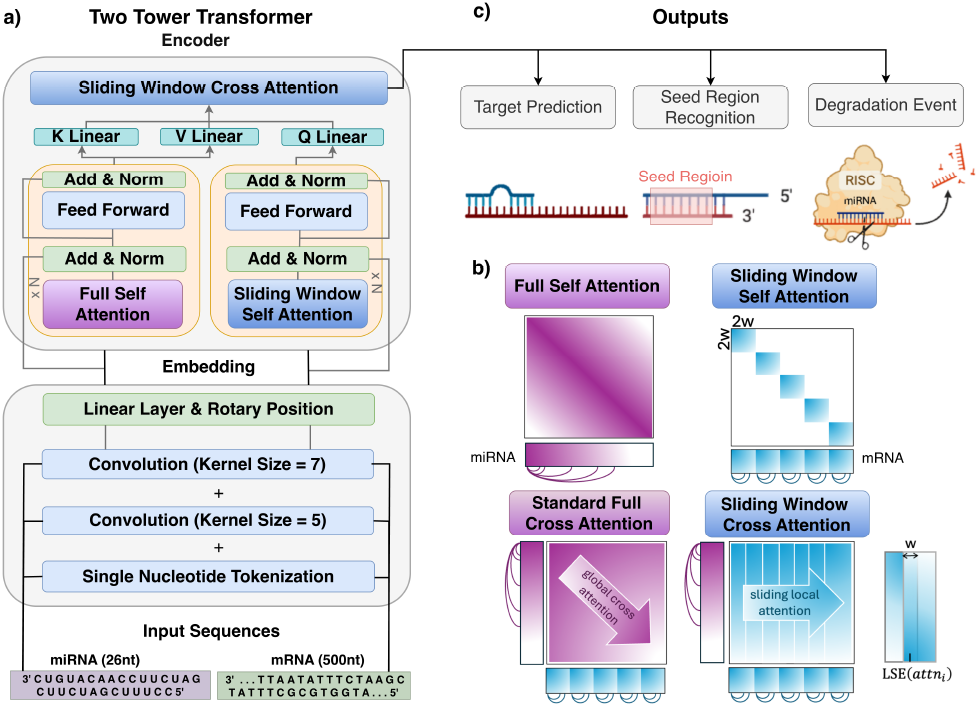
MiRformer overview. a) MiRformer leverages convolutional tokenization layers to extract seqeunce features from miRNA and mRNA, and encodes the features via a Dual-Transformer-Encoder Architecture (DTEA). The resulting embeddings are then fed into two dedicated transformer encoders for process miRNA and mRNA. The two embeddings are then fused via cross-attention that emulates the mRNA-mRNA nucleotide-level sequence recognition. b) Attention mechanisms. Full self attention encodes miRNA sequences by attending every base of the input miRNA sequence to itself and other bases. Sliding-window attention encodes mRNA sequences by attending only bases within a 2*w* window. Cross attention fuses both miRNA and mRNA sequence embeddings by scanning a 2*w* window across the mRNA embedding. c) Training tasks: target prediction, seed region recognition and degradation event prediction.

#### Dual-transformer-encoder architecture with distinct self-attention mechanisms

miRFormer uses two dedicated transformer encoders that are jointly trained end-to-end (Fig. 1a) with distinct attention mechanisms: **full self-attention** and **sliding-window attention** for miRNA and mRNA, respectively (Fig. 1b). Full self-attention encodes the complete miRNA sequence, allowing each nucleotide to attend to all other nucleotides. For mRNA sequences, we use sliding-window attention because it not only significantly reduces computational cost on modeling long sequences but also closely aligns with the biological mechanism of miRNA regulation where part of the 22-nt miRNA binds to small segment of a much longer mRNA sequence. Inspired by Longformer [Beltagy et al., 2020], sliding-window attention encodes the mRNA sequence using a window of size 2*w* with *w* = 20nt that slides across the mRNA sequence. The window of 2*w* width is centered on every nucleotide of mRNA sequence, allowing it to only attend to nucleotides inside the window. We segment the query and key mRNA embedding into overlapping chunks of 2*w*, compute attention within chunks, and merge chunks into the attention matrix without duplicate chunks. All matrix calculation is fully vectorized to reduce the time complexity from *O*(*L*^2^) with standard transformer attention to *O*(*wL*) for *L* mRNA length.

#### Cross-attention

Once we have the miRNA embedding and mRNA embedding, we need to design a sliding-window cross-attention mechanism to let miRNA sequence efficiently attend mRNA sequence, thereby mimicking the biological miRNA-mRNA ribonucleotide-level hybridization. We note that self-attention is a square matrix whereas cross-attention between miRNA and mRNA is a rectangular matrix, which requires a different design to merge the overlapping sliding windows (Fig. 1b). Let 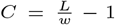 be the number of sliding-window chunks for an mRNA of length *L* with window width *w* = 20 nt. Each chunk is indexed by *c* ∈ {1, …, *C*} and each position inside a chunk is indexed by *i* = *c* × *w* + *u*, where *u* is the within-chunk offset. The overlapping length between adjacent chunks is 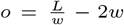. Before chunking, the cross-attention matrix is 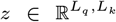 after merging overlapping windows, it becomes 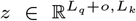 . To merge signals from all chunks *C*(*i*) covering the same position *i*, we apply LSE pooling: 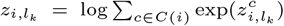 . We use LSE pooling instead of mean pooling because LSE preserves strong local alignment signals by approximating a max operator, while still aggregating information across windows. This prevents true binding-site signals from being diluted by low-attention regions, which frequently occurs when merging overlapping windows in long mRNA sequences. Finally, attention is normalized either along miRNA positions 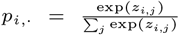 or along mRNA positions 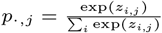(Fig. 5).

### 2.2 Datasets

#### TargetScan dataset

Positive samples were constructed from TargetScan (version 7.2) predictions with Context++ scores above a specified threshold (top 20% by percentile). For each positive miRNA-mRNA pair, we extracted a window around the seed region. Negative samples. To reduce degree/composition bias introduced by global shuffling, we constructed negatives in a miRNA-stratified, degree-preserving manner. For each miRNA, we sampled an equal number of negative mRNAs, where a negative mRNA is defined as an mRNA not predicted as a target of that miRNA by TargetScan. This design preserves the miRNA marginal frequency between positive and negative pairs and reduces the possibility that the model memorizes heterogeneous miRNA interaction degrees. Given a negative (miRNA, mRNA) pair, we then extracted seed-absent mRNA segments from the full 3’UTR sequence. If the 3’UTR length was shorter than the window length, we retained the full sequence. Otherwise, we sampled up to *N* windows of the specified length such that no window contains a ≥ 6-nt consecutive complementary seed match to the paired miRNA. We set *N* = 1 for 30-nt windows and *N* = 8 for 500-nt windows, with *N* ≤ *L* − *w* + 1 where *L* is the 3’UTR length and *w* is the window length. To investigate potential nucleotide composition bias in our negative sampling, we compared G/C and A/U content distributions between positive and negative mRNA sequences in the TargetScan 500 nt training set (Supplementary Section S2). The distributions overlap substantially, indicating that our negative sampling strategy does not introduce systematic nucleotide composition bias between classes. Overall, we merged human and mouse miRNA–mRNA pairs to build a dataset of 580,088 samples, consisting of 285,573 positive pairs with confirmed miRNA seed regions and 294,512 negative pairs without seed regions.

#### Seed position randomization

To prevent model from memorizing the location of seed regions, seed regions were randomly positioned within the 30-nt and 500-nt segment. The left extension was randomly chosen between total extension and seed start, where total extension = window length - seed length. The right extension is total extension - left extension. If the right extension exceeds the mRNA sequence length, the window is left-adjusted accordingly. Similarly for the left extension.

We constructed a dataset that includes miRNA IDs, mRNA IDs, miRNA sequences, mRNA windows, an interaction label (with positive samples annotated as 1 and negative samples as 0), as well as the seed start and seed end positions within each mRNA window. For negative samples, the seed positions were set to −1 and were excluded from model training. The data were divided into separate training (n=522,077), validation (n=46,408), and test sets (n=11,603). Only training and validation sets were seen during model training.

#### Degradome-seq dataset

The Human Degradome-seq data consists of experimentally observed cleavage sites from PARE and GMUCT datasets [Liu et al., 2015], retrieved from the Gene Expression Omnibus (GEO) and Sequence Read Archive (SRA) via StarScan. Cleavage loci annotations were obtained by asking authors of starBase [Li et al., 2014a]. Sequence of reference genome and gene annotation file can be downloaded from GENCODE v19 for assembly hg19 at https://www.gencodegenes.org/human/release_19.html. The 3’UTR sequences were derived from GENCODE v19 annotations [Harrow et al., 2012] for the GRCh37 (hg19): `gffread gencode.v19.annotation.gtf -g GRCh37.p13.genome.fa -w transcripts.fa`. Cleavage site genomic coordinates were subsequently converted to 3’UTR transcript coordinates using the same GENCODE v19 reference. The length of the longest sequence reached 25k nucleotides. Next, we ensured the quality of the dataset by 1) filtering for cleavage sites in 3’UTR, 2) filtering for sites observed in at least 2 experiments and 3) filtering for sites having a false discovery rate (FDR) less than 0.01. After filtering, 13,084 high-quality cleavage sites were retained for model training and evaluation. Next, we constructed a 500-nt dataset by sampling 500-nt windows containing the cleavage sites. Both the full-length 3’UTR and the 500-nt window datasets were partitioned into training (n=10,467), validation (n=2094), and test sets (n=525). Only training and validation sets were seen during model training.

#### Paired Human Degradome-seq with TargetScan Human

We emphasize that throughout our study, we used the filtered 13,084 high quality Degradome-seq dataset for cleavage site prediction training and evaluation. Nevertheless, we investigated co-localization of TargetScan-predicted seed and annotated Degradome-seq cleavage sites (Supplementary Section S3). Sparse matching pairs of TargetScan and Degradome-seq data were identified by mapping genomic coordinates of cleavage sites to trancriptomic coordinates of TargetScan 3’UTR blocks. Due to the indenpendent nature of the two datasets, and our strict critera in matching mRNA-miRNA pairs of the two datasets (Supplementary Section S3), we decided to use the filtered experimental Degradome-seq dataset for training and evaluation, rather than the paired subset, to maximize both the data quality and the statistical power of our analyses.

#### miRBench datasets

To evaluate generalizability to independent experimental data, we used the miRBench benchmark [Sammut et al., 2025], which provides bias-corrected miRNA binding site interaction datasets derived from CLASH and chimeric eCLIP experiments. miRBench addresses the miRNA frequency class bias present in existing datasets by ensuring that miRNA family frequencies are balanced between positive and negative classes. We used four test sets: Klimentova2022 test (477 positives, chimeric eCLIP), Hejret2023 test (495 positives, CLASH), Manakov2022 test (168,342 positives, chimeric eCLIP), and Manakov2022 left-out (10,027 positives, containing miRNA families unique to this dataset). For retraining experiments, we used the Hejret2023 train set (4,084 positives) and Manakov2022 train set (1,253,320 positives). All datasets are class-balanced (1:1 positive to negative ratio). Binding site sequences are short (50 nt) chimeric read fragments, which we processed using our 30 nt model configuration.

### 2.3. Prediction Tasks

The 3 tasks are illustrated in Fig. 1c and described below.

#### Target prediction

Given a pair of miRNA and mRNA, MiRformer was trained to classify each miRNA–mRNA pair as interacting (positive) or non-interacting (negative). Specifically, the output embeddings from the last cross-attention layer were fed into a 3-layer multi-layer perceptron (MLP), i.e. 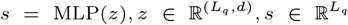 where *d* is the embedding dimension (Section 2.5). We explored three ways to get target predictions: 1. through mean-pooling, 2. through temperature-regulated LSE-pooling and 3. through a global CLS token. In mean pooling, logits are 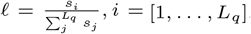. In LSE-pooling, logits are obtained by 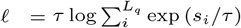,where *τ* is the temperature parameter. The smaller *τ* is, the closer it is to a max-pooling. The greater *τ* is, the closer it is to a mean-pooling. We tested *τ* = {0.5, 0.7, 1.0} for finetuning the temperature parameter. Through the global CLS token, the CLS token was appended to the beginning of every sequence and has global view of the entire sequence. It was isolated from the rest of the sequence and used to predict target. We later showed the effect of the three ways to get target predictions in Results 3.2.

#### Seed-region recognition

Given a pair of miRNA and target mRNA, MiRformer was trained to predict the start and end positions of the seed region within the mRNA sequence. The output embeddings from the last cross-attention layer are fed into a linear layer to give the nucleotide-level embedding 𝓁_*c*_, and is sequentially fed to softmax to give the probabilities at every nucleotide of either 1) being the start of a seed region or 2) being the end of a seed region. For both tasks, MiRformer was trained on TargetScan Human and Mouse v7.2 [Agarwal et al., 2015].

#### Degradation event prediction

Given a pair of miRNA and target mRNA, MiRformer was trained to predict cleavage sites coordinates within 500-nt windows and cleavage sites coordinates within full-length 3’UTR transcripts.

### 2.4 Objective functions

MiRformer minimizes a combined loss function for task 1 and 2 described above: ℒ = *α*_1_ℒ_1_ + *α*_2_ℒ_2_ where ℒ_1_ is the binary cross-entropy loss for target prediction:

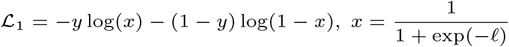

and 𝓁 is the sequence-level embedding; ℒ_2_ is the cross-entropy loss for seed-region recognition:

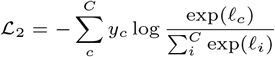

where 𝓁_*c*_ is the logit at nucleotide *c. C* is the total number of nucleotide of the input sequence. We set *α*_1_ = 1 and *α*_2_ = 0.75. Note that ℒ_2_ is computed only for positive samples.

After training on TargetScan data, MiRformer was continuously trained on Degradome-seq data to predict cleavage sites within mRNA sequences. We transformed cleavage is site coordinates into Gaussian-smoothed labels of size 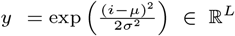, where *i* is the query position and *µ* the cleavage site position, *L* is mRNA length, and standard deviation *σ* = 3.0. We define the loss as a Gaussian-weighted binary cross entropy (BCE) loss. Let ℒ_*i*_ = −*y*_*i*_ log(*ŷ*) − (1 − *y*_*i*_) log(1 − *ŷ*_*i*_) and *w*_*i*_ = 1 + (*λ* − 1)*y*_*i*_ where *λ* = [1, 2] is a weighting factor to enhance the gradient at *y*_*i*_ = 1. The weighted BCE loss is 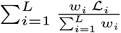.

### 2.5. Hyperparameter tuning

We optimized the model’s hyperparameters to identify the configuration that yielded the best overall performance —specifically, the set of values that minimized training loss and, on the validation set, achieved the highest binding accuracy and F1 score for overlap between the predicted seed match and the TargetScan seed match (Table 1).

**Table 1.**
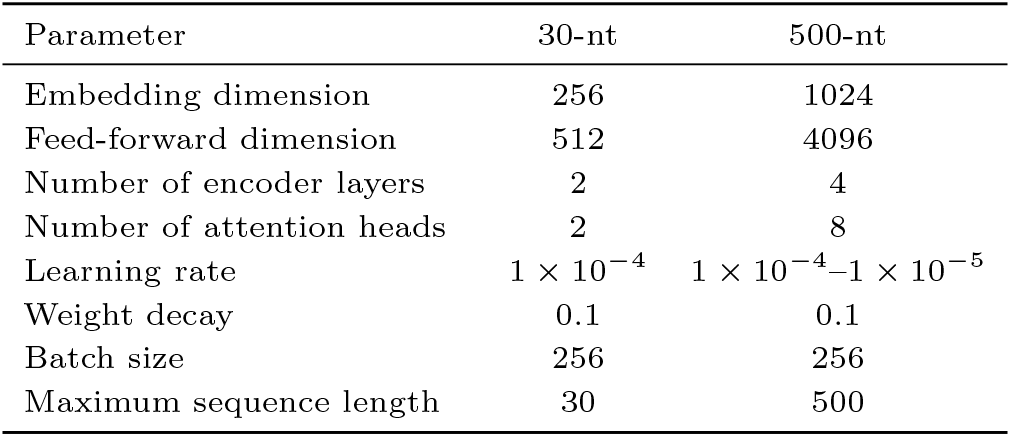
Optimal hyperparameters for MiRformer model.

| Parameter | 30-nt | 500-nt |
| --- | --- | --- |
| Embedding dimension | 256 | 1024 |
| Feed-forward dimension | 512 | 4096 |
| Number of encoder layers | 2 | 4 |
| Number of attention heads | 2 | 8 |
| Learning rate | $1 \times 10^{-4}$ | $1 \times 10^{-4} - 1 \times 10^{-5}$ |
| Weight decay | 0.1 | 0.1 |
| Batch size | 256 | 256 |
| Maximum sequence length | 30 | 500 |

### 2.6. In-silico mutagenesis

We perturbed every base in mRNA sequence, from 5’ to 3’ including both the seed and non-seed region, and recorded the change in probability, 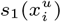, and the change in attention score, 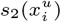. Specifically, each base was perturbed 3 times from the original base to the 3 alternative bases. We then averaged the change over the 3 perturbations. Formally, 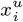 denotes the substituted base *u* at position 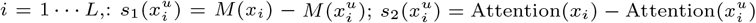, where *M* (*x*_*i*_) denotes the model’s prediction probability for the original sequence, and 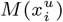 denotes the prediction probability after replacing base *x*_*i*_ with base *u*. The final perturbation score is 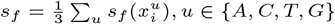.

### 2.7. Baseline methods

For miTAR, since it was only trained on 66 nt miRNA-mRNA concatenated length using 24-nt miRNA, we scanned the 500 nt mRNA by a 40-nt window and predicted seed regions within the window. For target prediction, we ran miTAR and scanned every 40-nt window and selected the window with the highest probability. For seed region prediction, we added a span head that was a copy of the BiLSTM inside the classification head and also scanned every 40-nt window for seed regions. The window having the highest sum of top-k (*k* = 8) probability was evaluated on seed start and end prediction. For REPRESS, the software requires the input sequence length to be 12.5k and pads the input to the same maximum length. Its outputs include probability of binding for 29 cell lines at each base pair as well as read counts of cleavage sites for 10 cells lines. We evaluated the outputs as described in [Kanuparthi et al., 2025] - average pooled both the prediction and labels within the predefined window lengths for each cell line and divided the output into windows. For Mimosa, we slided a 40-nt window across each mRNA, scoring each window with the pretrained Mimosa Transformer, and classify the miRNA-mRNA as interactive if the best window’s probability exceeds a threshold. We predicted seed start and end by inferring the paired region within the best-scoring window (via the same seed-alignment pairing map used by Mimosa) and aggregated per-nucleotide seed-span scores across windows. In a similar manner, we predicted cleavage sites by identifying the cleavage position within the best-scoring window based on seed alignment and mapping the inferred site back to the original mRNA sequence.

## 3. Results

### 3.1. Improved performance and interpretability

As a proof-of-concept, we first evaluated the benefits of having the hybrid tokenization on short 30-nt and compared its performance with the one without convolution layer (Fig. 2). Model trained with convolutional tokenization demonstrated better performance than the ablated counterparts (Fig. 2a). We investigated the model’s interpretability by showing cross attention scores with and without convolution layers. We observed that ablated model attended to the boundaries of seed region instead of the nucleotides inside the seed (Fig 2b). In contrast, MiRformer with the convolution layer produced more prominent cross attention score at not only the boundaries but also the nucleotides inside the seed region (Fig. 2c).

**Fig. 2.**
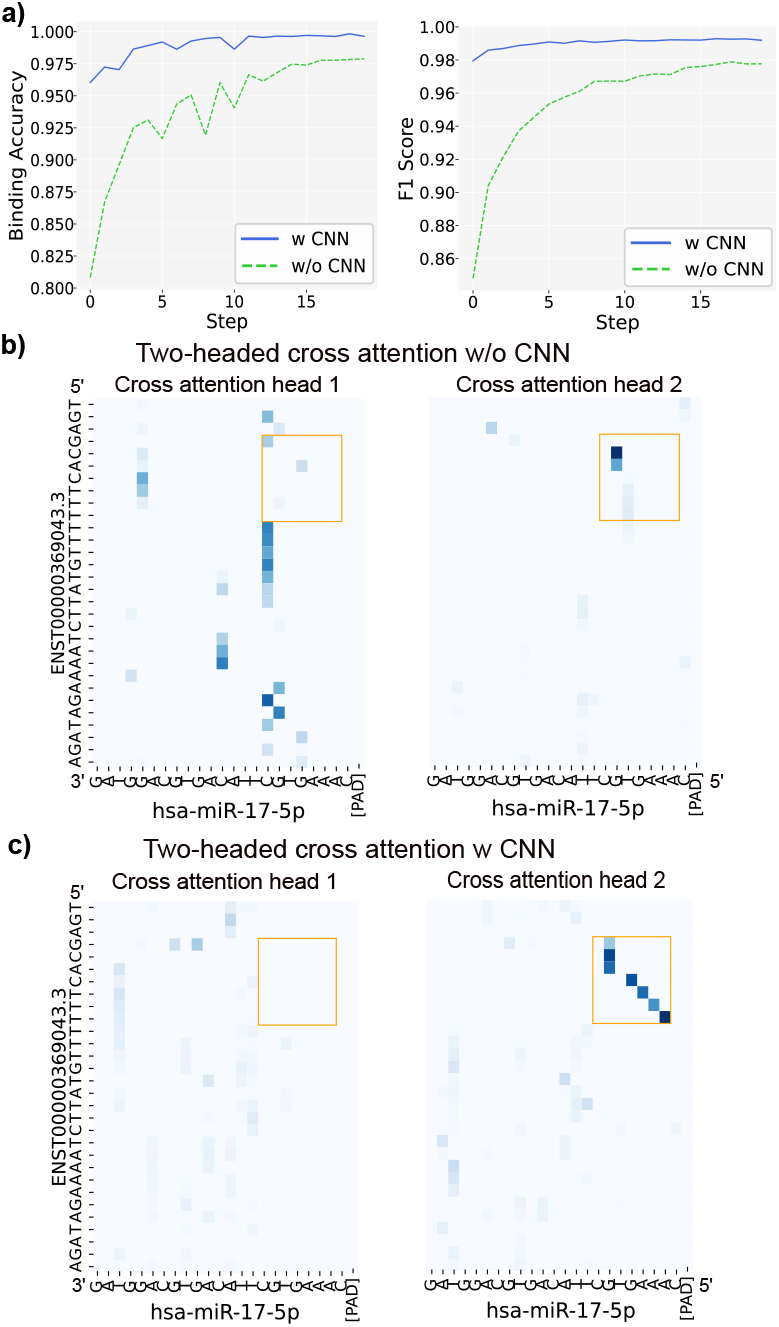
Proof-of-concept analysis on 30-nt short mRNA sequences. a) Model performance comparison between with and without convolutional feature extraction. b) Heatmap produced by model trained without convolution feature embeddings. c) Heatmap produced by model with convolution feature embeddings. For both b and c, cross attention weights from two attention heads are shown as heatmaps produced by a dual-transformer-encoder architecture trained on both binding labels and seed span positions. Complimentary seed region within mRNA (y-axis) and miRNA (x-axis) are highlighted in box.

To further investigate model’s interpretability on single-base level, we performed in-silico mutagenesis (ISM) and averaged probabilities at every base of all perturbations (Section 2.6). We showed change in target prediction scores and change in cross attention scores after ISM for randomly selected examples. The changes were the highest within seed regions (Fig. 3a-c). This provides evidence that our model predicts miRNA-mRNA bindings by identifying the seed regions. We also tested model’s ability on predicting targets on negative miRNA-mRNA pairs with artifically inserted seed regions (Fig. 3d). We observed a significant difference in target prediction scores between the original negative miRNA-mRNA pairs and the same negative pairs after insertion of seed regions. This further shows that our model is able to predict miRNA targets based on true seed regions. Overall, convolutional layers enable effective local feature extraction that improves both predictive performance and attention interpretability.

**Fig. 3.**
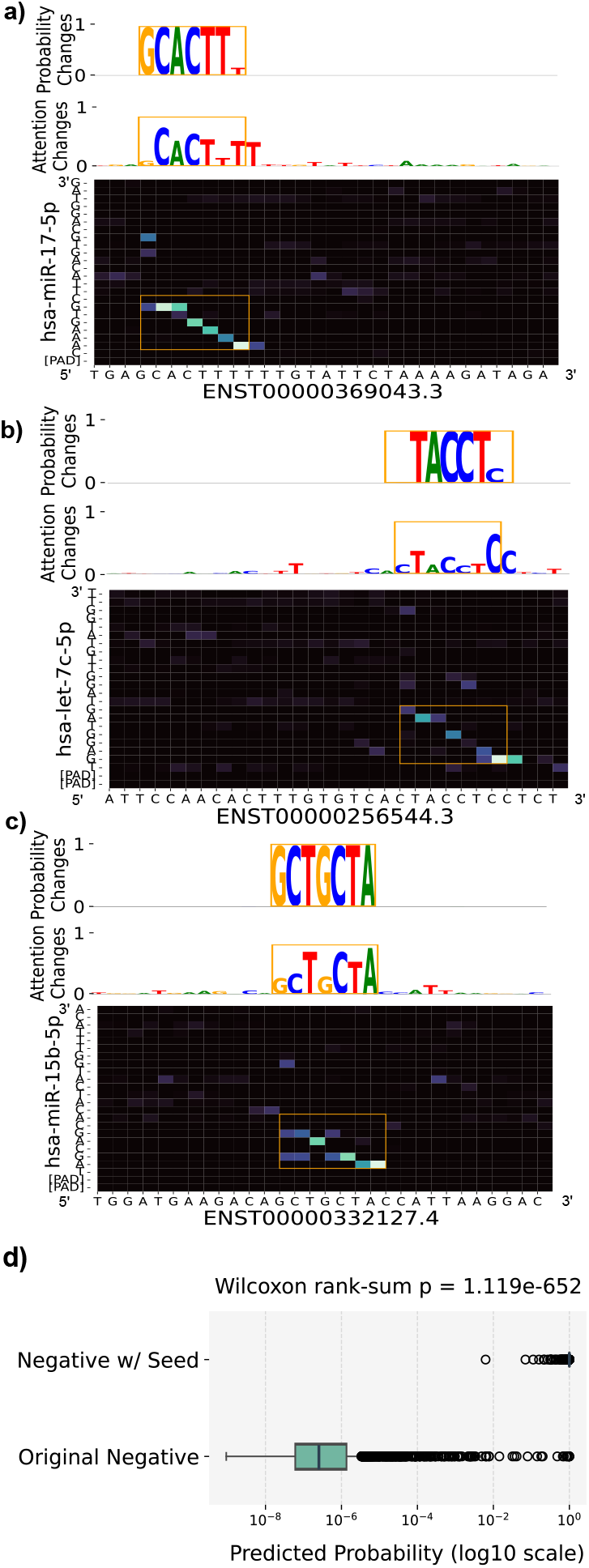
In-Silico Mutagenesis (ISM) analysis of 30-nt mRNA sequences. (a-c) ISM and attention visualization. We perturbed every base and recorded the change in target prediction probability and the changes in cross attention matrix at each base. For each of the 3 examples, the top two tracks display the probability and attention change at each base. The bottom heatmap displays the cross-attention between miRNA and mRNA. d) Boxplot showing predicted interaction probabilities on Original Negative – a set of negative miRNA-mRNA pairs without seeds, and Negative w/ Seed - Original Negative after inserting seed regions.

### 3.2. Modeling long mRNA sequences

We evaluated the proposed sliding-window cross-attention and LSE (Log-Sum-Exponential) pooling on 500-nt mRNA sequences by comparing 7 model variants with different combinations of pooling methods, LSE temperature values (*τ* = 0.5, 0.7, 1.0), and a CLS-token-only baseline for target prediction (Section 2.3). All variants were monitored on a held-out validation set sampled from the TargetScan dataset (Section 2.2, Fig. 4).

**Fig. 4.**
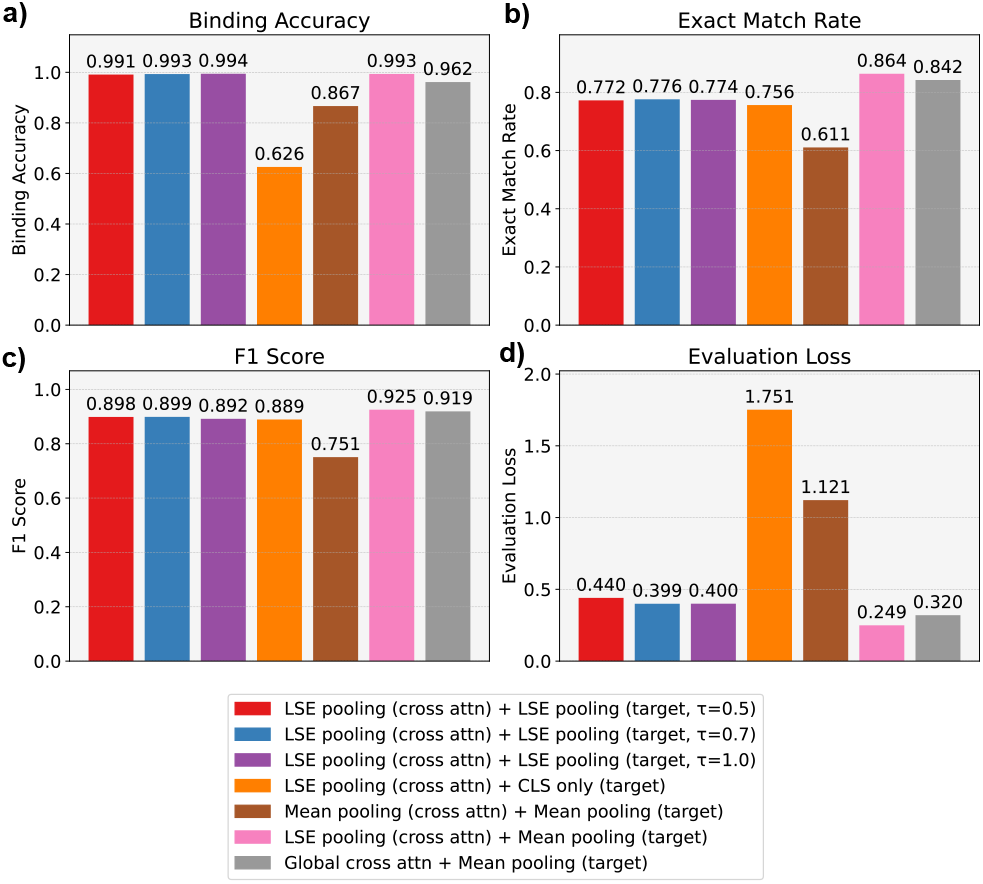
Pooling methods comparison. The pooling method is followed by the model layer to which it is applied (for example, LSE pooling (cross attn) indicates that LogSumExp pooling is used in the cross-attention layer, while Mean pooling (target) indicates that mean pooling is used in the target prediction head). *τ* denotes the temperature parameter used in LogSumExp pooling. a) Accuracy of the predicted miRNA–mRNA interactions. b) Accuracy of the predicted seed start and seed end positions. c) F1 score quantifying the overlap between predicted seed regions and the true seed regions. d) Loss for each method, evaluated on an independent validation set for both target and seed predictions.

LSE pooling consistently outperformed mean pooling across all metrics. Most notably, LSE improved seed-region F1 score by ∼0.20 over mean pooling (Fig. 4c), supporting our hypothesis that LSE successfully amplifies sparse seed signals rather than diluting them through averaging during window aggregation. Models with LSE at the cross-attention layer also outperformed the global cross-attention variant on both binding accuracy (0.993 vs. 0.962; Fig. 4a) and seed-region F1 (0.925 vs. 0.919; Fig. 4c). Performance was stable across temperature values *τ* = 0.5, 0.7, 1.0, with *<*2% variation on all metrics, suggesting that *τ* does not require task- or species-specific tuning. Using only the CLS token for target prediction reduced accuracy by 0.40 (Fig. 4a), confirming that a single summary token cannot capture sufficient information from long mRNA contexts.

### 3.3. Attention interpretability

To verify that our model truly learned seed regions in 500-nt mRNA sequence context, we examined the final cross-attention scores from the sliding-window cross-attention layer in similar way as for the 30-nt analysis (Section 3.1). We also conducted ISM on mRNA sequence to investigated the importance of each base (Section 2.6). In a selected example, we observed that miRformer focused on seed regions both in cross-attention scores and signals from ISM tests (Fig. 5). We tried two ways to normalize cross attention: one was normalizing by mRNA length (Fig. 5a) and the other was normalizing by miRNA length (Fig. 5b). Normalizing by mRNA (miRNA) length was analogous to inquiring which ribonucleotides inside the mRNA (miRNA) sequence does the miRNA (mRNA) attends to the most. We observed much clearer pattern at the boundaries of the seed region when normalizing by miRNA (Fig. 5b). Nevertherless, attention heatmap pattern is much clearer in seed region compared to non-seed region when normalizing by mRNA (Fig. 5a), which underscored the importance of our sliding-window attention and LSE pooling on longer sequences. In contrast, mean pooling produces much noisier signals (Fig. 5c). Although global attention produced high prediction performance (Fig. 4), the resulting attention is not interpretable (Fig. 5d). In contrast, we observed a significant difference in average cross-attention scores between seed and non-seed regions for all sliding window approaches (Fig. 5a-c). However, the method of pooling proved critical. While the cross attention + mean probability approach (Fig. 5c) did yield a significant difference, the statistical separation was orders of magnitude lower (*p* ≈ 10^−16^) compared to the cross attention + LSE approaches (*p* ≈ 10^−49^ to *p* ≈ 10^−67^). This suggests that mean pooling dilutes the strong signals from seed regions by averaging them with the surrounding noise. Therefore, MiRformer utilizing sliding window cross-attention with LSE pooling provides the optimal balance - conferring highly interpretable attention without compromising performance.

**Fig. 5.**
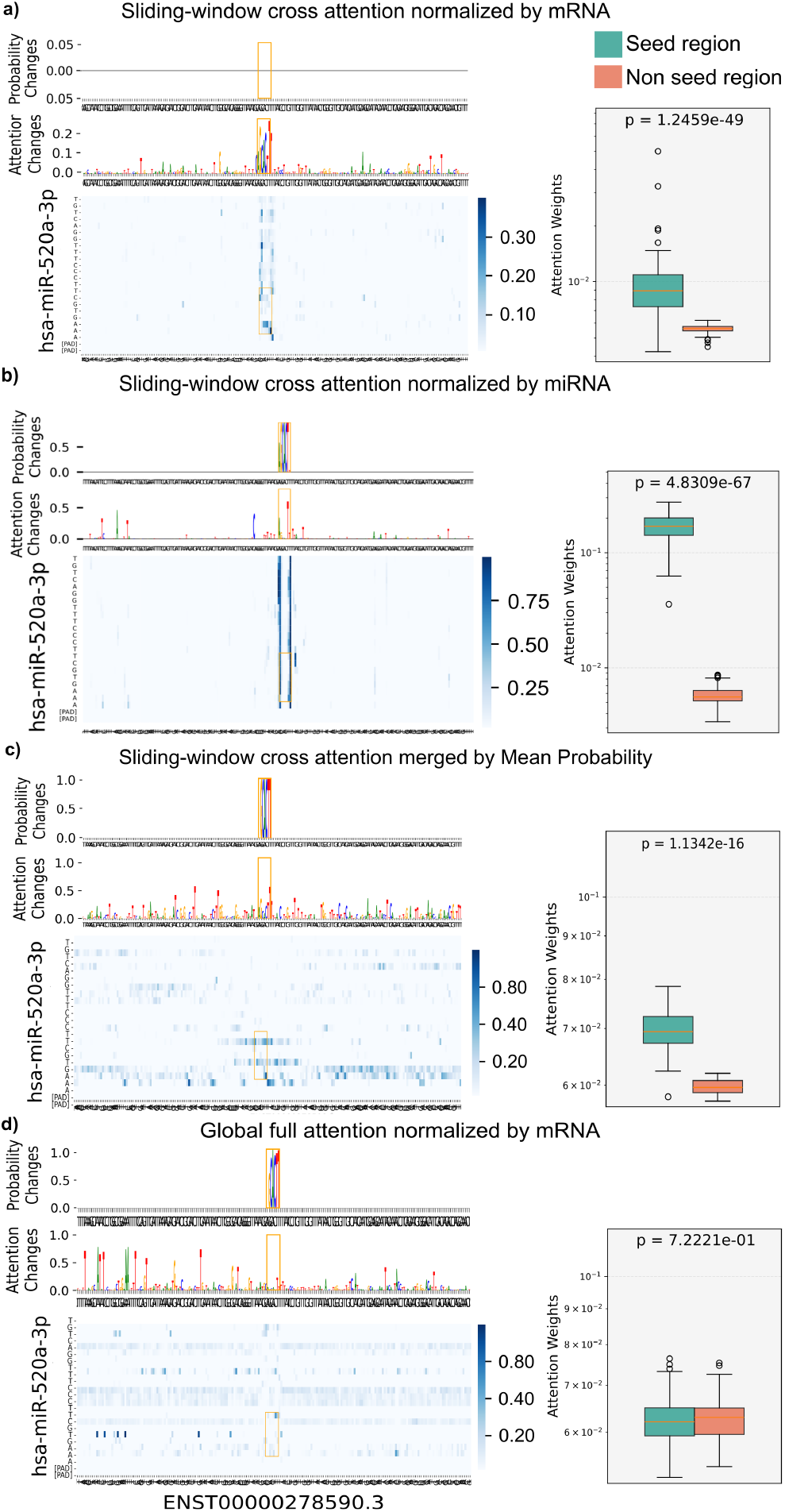
Model interpretability through cross attention heatmaps on a 500-nt mRNA sequence. For each panel, top track displays the change in probability at the single nucleotide resolution after ISM; middle track shows the change in attention after ISM; bottom track displays cross attention between miRNA and mRNA at the nucleotide resolution without ISM perturbation. Boxplots show the average attention weights distributions of seed and non-seed regions. P-values were calculated by Wilcoxon rank-sum test on 200 miRNA-mRNA pairs between base pairs in the seed region and base pairs outside the seed region. a) Sliding-window cross-attention + LSE pooling and normalized by mRNA. b) Sliding-window cross attention + LSE pooling and normalized by miRNA.c) Sliding-window cross attention + mean pooling and normalized by mRNA. d) Global cross attention and normalized by mRNA.

### 3.4. MiRformer performs well on all tasks

We evaluated MiRformer against three state-of-the-art models (REPRESS, miTAR and Mimosa) for 3 tasks - target prediction, seed region recognition and experimental degradation event prediction (Section 2.3) and measured all models performance by 6 metrics (Fig. 6). For the first two tasks, we used the TargetScan Human and Mouse (version 7.2) as labels, which were directly downloaded from the TargetScan database. All methods were tested on 500-nt mRNA 3’UTR segments. MiRformer outperformed other models on all 6 metrics (Fig. 6 top row). Notably, MiRformer exceeded the second best model on predicting Degradome-seq cleavage sites for Hit at 5-nt accuracy by 0.348 and on Hit at 3-nt by 0.323.

**Fig. 6.**
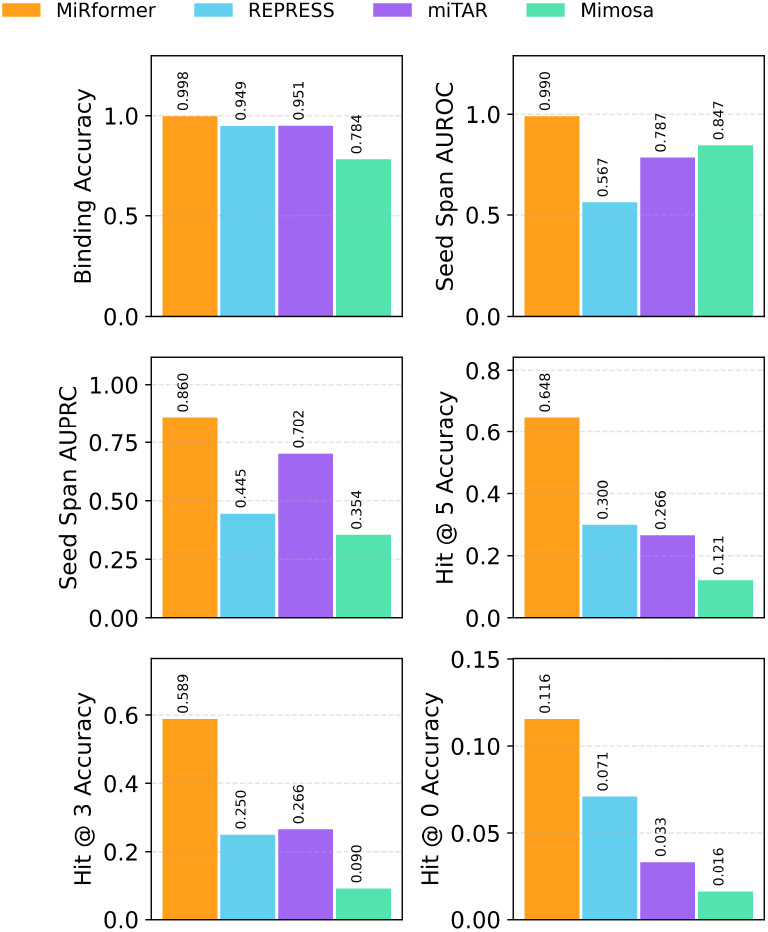
Benchmark on seed region and cleavage site prediction. Binding Accuracy evaluates classification of miRNA-mRNA interactions. Seed Span AUROC and AUPRC evaluate nucleotide-level prediction for inside the seed region. Hit @ k Accuracy measure accuracies of predicting the cleavage site within k-nt distance to the true cleavage site in the target mRNA sequence.

### 3.5. Generalizability to miRBench data

To assess whether MiRformer captures genuine biological interaction patterns beyond TargetScan’s heuristic rules, we evaluated on miRBench [Sammut et al., 2025], an independent benchmark derived from CLASH and chimeric eCLIP experimental data with bias-corrected negative sampling. Critically, miRBench negatives retain seed-complementary regions, so a model relying solely on seed presence would perform at chance level. We conducted three evaluations (Supplementary Table S1): (1) zero-shot evaluation of our TargetScan-trained model, (2) training on the Hejret2023 train set (4,084 positive pairs), and (3) training on the Manakov2022 train set (1.25M positive pairs). In the zero-shot setting, MiRformer achieves APS of 0.73–0.77 across all four test sets, competitive with state-of-the-art models trained directly on experimental data (e.g., TargetScanCNN: 0.71– 0.77; miRBind: 0.71–0.80). This demonstrates that MiRformer learned generalizable biological patterns from TargetScan data rather than memorizing TargetScan’s rules. When retrained on the large Manakov2022 dataset, MiRformer achieves APS of 0.82–0.83 on three test sets, comparable to the best results reported in the miRBench benchmark. Performance on the Manakov2022 left-out set—containing miRNA families entirely absent during training—was lower (APS 0.70), a limitation that could be addressed with more diverse training data (Supplementary Table S1).

### 3.6. Computational scalability

We profiled MiRformer’s inference time and peak GPU memory across mRNA lengths from 500 to 25,000 nt comparing three configurations: MiRformer with sliding-window attention, a variant using standard full attention, and RNAhybrid [Rehmsmeier et al., 2004] (Supplementary Section S1). Both inference time and GPU memory scale linearly with sequence length for MiRformer’s sliding-window attention, confirming the theoretical *O*(*wL*) complexity. In contrast, the full attention variant exhibits quadratic growth and exceeds 24GB GPU memory at 14,000nt. RNAhybrid, a lightweight CPU-based dynamic programming tool, achieves faster per-pair inference with minimal memory usage ∼34 MB(Supplementary Figure S1). These results demonstrate that MiRformer enables efficiently process full-length 3’UTRs and long non-coding RNAs on a single GPU.

### 3.7. Jointly inferring miRNA-binding & cleavage sites

While Degradome-seq provides experimental evidence on miRNA-mRNA interactions via observed cleavage sites, it does not produce the exact miRNA binding sites. TargetScan generates computational seed-match predictions, but does not predict AGO2 cleavage sites.

We sought an approach to simultaneously predict both seed region span and cleavage sites. Specifically, we first trained MiRformer on TargetScan data and then continuously trained it on Degradome-seq data for predicting cleavage sites (Section 2.4). We tested MiRformer on two datasets: 1. Test 500 - a TargetScan subset containing 500-nt mRNA paired with miRNA unseen during training. 2. UTR - complete full-length 3’UTR of mRNAs of up to 25,000 nt (Fig. 7).

**Fig. 7.**
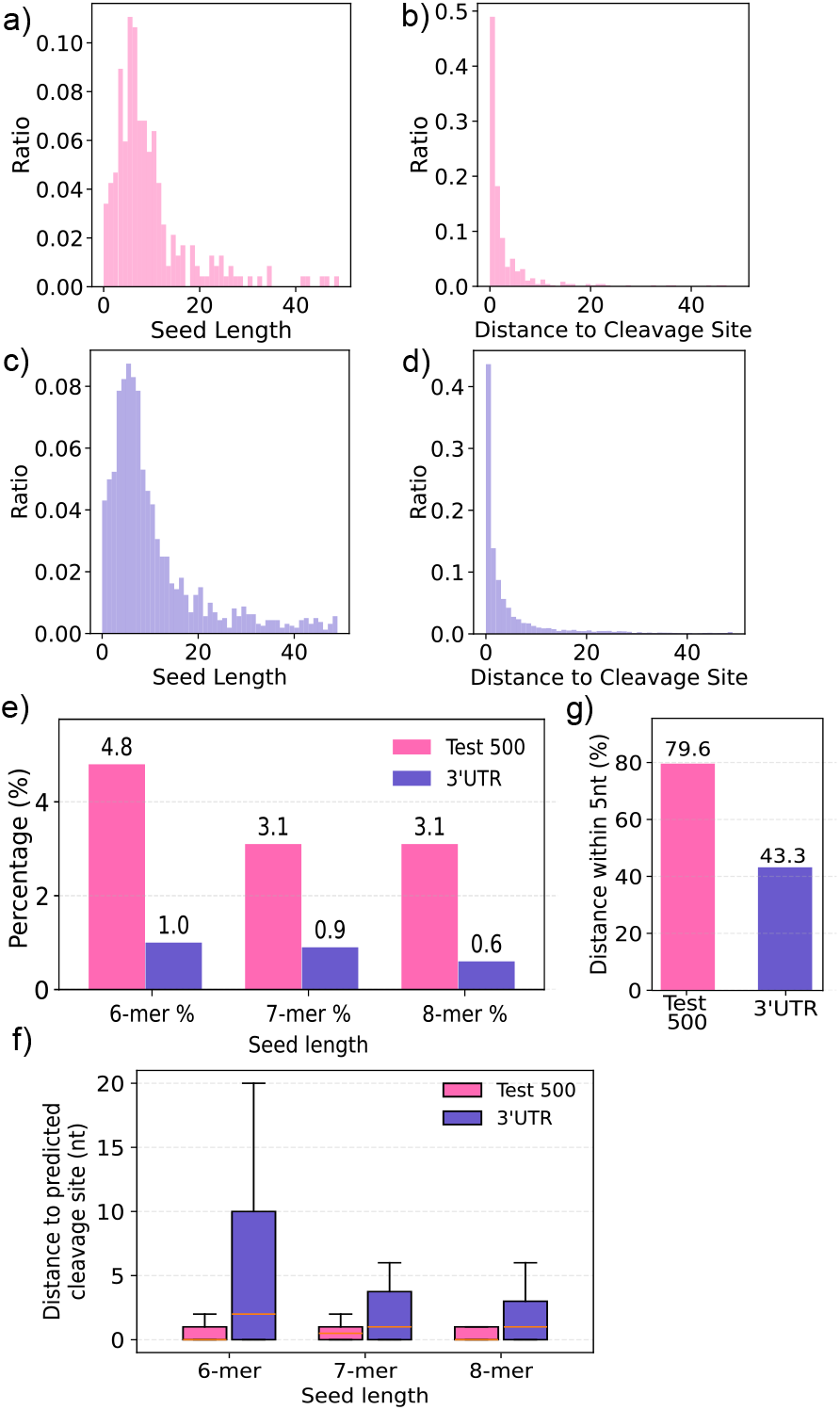
Predicted seed regions and cleavage sites. a) Length distribution of the predicted seed regions on the Test 500 dataset (i.e., 500-nt mRNA 3’UTR segments). b) Distribution of the distance between the predicted seed regions and the predicted cleavage sites on Test 500 dataset. c,d) Same as a and d but for the predictions on the full-length 3’UTRs. e) Proportion of predicted canonical seed lengths (6-, 7-, and 8-mers) in Test 500-nt and full-length 3’UTR sequences. f) Distribution of distance between predicted canonical seed lengths (6-, 7- and 8-mers) and the predicted cleavage sites. g) Proportion of predicted cleavage sites that fall within 5-nt to seed regions for 500-nt and 3’UTRs.

For the <u>500-nt mRNA segments</u>, most of the predicted seed regions span 0 to 10 nt (Fig. 7a) and fell within 0 to 10 nt distance from the predicted cleavage sites (Fig. 7b). About 4.8% of them were 6-mer, 3.1% were 7-mer and 3.1% were 8-mer (Fig. 7e). Also, most of the predicted 6, 7 or 8-mer seed regions fell within only 2.5 nt from the predicted cleavage sites (Fig. 7f).

For the <u>full-length 3’UTR sequences</u>, 3’UTR sequences were padded to the longest in a one batch to predict both the seed regions and the cleavage sites. Most of the predicted seed regions were within 0 to 20 nt (Fig. 7c) and located within 10 nt from the predicted cleavage sites (Fig. 7d). Over 1.0% of all predicted seed regions were 6-mers, 0.9% were 7-mer and 0.6% were 8-mer (Fig. 7e). Lastly, 79.6% and 43.4% of predicted cleavage sites were within 5 nt distance to seed regions within 500-nt and full-length 3’UTR, respectively (Fig. 7g). Together, we demonstrated that MiRformer was capable of simultaneously predicting seed regions and cleavage sites despite not being previously trained such paired data.

## 4. Discussion

Characterizing sequence features governing the miRNA–mRNA interactome remains a major challenge [Li and Zhang, 2015]. Traditional methods such as TargetScan rely on handcrafted features (e.g., seed matching and conservation) but miss many exceptions, while CNN-based models struggle to capture nucleotide-level interactions over long contexts. Transformer-based models offer a solution by modeling both local and global dependencies via attention. Here, we introduce MiRformer, a dual-encoder transformer that jointly models miRNA and mRNA sequences using sliding-window cross-attention with LSE pooling. This design scales to long sequences and yields interpretable attention patterns. Across benchmarks, MiRformer delivers accurate and interpretable predictions for binding, seed localization, and cleavage-site identification.

MiRformer is primarily trained on TargetScan-derived labels, which rely on heuristic rules such as canonical seed matching and conservation. To assess potential bias toward these rules, we evaluated the model on miRBench [Sammut et al., 2025], an independent benchmark based on experimental CLASH and chimeric eCLIP data with seed-matched negatives. Without retraining, MiRformer achieves competitive performance (APS 0.73–0.77) across all test sets, indicating it captures interaction patterns beyond seed presence. When retrained on experimental eCLIP data (Manakov2022), it matches the best reported miRBench results, demonstrating robustness across data sources. Although we demonstrated MiRformer generalizes well to miRBench datasets, that contains nearly 40% non-canonical interactions, training on TargetScan may still limit generalization to non-canonical binding sites. In addition, our negative sampling strategy cannot guarantee that TargetScan-absent pairs are true negatives, introducing potential label noise. Performance drops on the Manakov2022 left-out set (APS 0.70), suggesting limited generalization to unseen miRNA families. Future work should incorporate larger experimental datasets, leverage cross-species alignments, and fine-grained hyperparameter tuning.

Looking ahead, MiRformer could serve as a foundation to learn RNA structural context from sequence via multimodal transformers, eliminating reliance on heuristic features such as minimum energy. It can be extended to incorporate single-cell expression profiles [Hücker et al., 2021], and miRNA synergistic network [Li et al., 2014b, Liu et al., 2024] to further generalize to non-canonical binding sites. Pretrained RNA language models across diverse transcriptomes could further enable zero-shot inference for novel miRNAs and disease variants. Together, these directions shift miRNA modeling from feature engineering to data-driven reasoning, supporting discovery of new regulatory functions and RNA therapeutics. We view MiRformer as a step toward general, data-driven modeling of post-transcriptional regulation.

## 5. Author contributions

Y.L. conceived the study and supervised the project. J.G. implemented the method and conducted the experiments. All authors wrote and reviewed the manuscript.

## 6. Acknowledgments

J.G. was supported by DNA2RNA Scholarship in promoting innovative RNA therapies. Y.L. is supported by CRC-2021-00547 and CIHR PJT-540722. Special thanks to Keren Zhou who kindly provided access to Degradome sequencing data.

## References

V. Agarwal, G. W. Bell, J.-W. Nam, and D. P. Bartel. Predicting effective microrna target sites in mammalian mrnas. elife, 4:e05005, 2015.

Ž. Avsec, V. Agarwal, D. Visentin, J. R. Ledsam, A. Grabska-Barwinska, K. R. Taylor, Y. Assael, J. Jumper, P. Kohli, and D. R. Kelley. Effective gene expression prediction from sequence by integrating long-range interactions. Nature Methods, 18(10):1196–1203, 2021.

D. P. Bartel. Micrornas: genomics, biogenesis, mechanism, and function. cell, 116(2):281–297, 2004.

Beltagy, M. E. Peters, and A. Cohan. Longformer: The long-document transformer. 2004.05150, 2020.

Y. Bi, F. Li, C. Wang, T. Pan, C. Davidovich, G. I. Webb, and J. Song. Advancing microRNA target site prediction with transformer and base-pairing patterns. Nucleic Acids Research, 52(19):11455–11465, 2024.

H. Dalla-Torre, L. Gonzalez, J. Mendoza-Revilla, N. Lopez Carranza, A. H. Grzywaczewski, F. Oteri, C. Dallago, E. Trop, B. P. de Almeida, H. Sirelkhatim, G. Richard, M. Skwark, K. Beguir, M. Lopez, and T. Pierrot. Nucleotide transformer: building and evaluating robust foundation models for human genomics. Nature Methods, 22(2):287–297, 2024.

T. Gu, X. Zhao, W. B. Barbazuk, and J.-H. Lee. mitar: a hybrid deep learning-based approach for predicting mirna targets. BMC bioinformatics, 22:1–16, 2021.

J. Harrow, A. Frankish, J. M. Gonzalez, E. Tapanari, M. Diekhans, F. Kokocinski, B. L. Aken, D. Barrell, A. Zadissa, S. Searle, et al. Gencode: the reference human genome annotation for the encode project. Genome research, 22(9):1760–1774, 2012.

V. Hejret, N. M. Varadarajan, E. Klimentova, K. Gresova, I.-C. Giassa, S. Vanacova, and P. Alexiou. Analysis of chimeric reads characterises the diverse targetome of AGO2-mediated regulation. Scientific Reports, 13(1):22895, 2023. ISSN 2045-2322.

S. M. Hücker, T. Fehlmann, C. Werno, K. Weidele, F. Lüke, A. Schlenska-Lange, C. A. Klein, A. Keller, and S. Kirsch. Single-cell microrna sequencing method comparison and application to cell lines and circulating lung tumor cells. Nature communications, 12(1):4316, 2021.

Kanuparthi, S. E. Pour, S. D. Findlay, O. Wagih, J. M. Gutierrez, R. Gao, J. Wintersinger, J. Lin, M. Gabra, E. Bohn, et al. Sequence based prediction of cell type specific microrna binding and mrna degradation for therapeutic discovery. bioRxiv, pages 2025–05, 2025.

J.-H. Li, S. Liu, H. Zhou, L.-H. Qu, and J.-H. Yang. starBase v2. 0: decoding mirna-cerna, mirna-ncrna and protein–rna interaction networks from large-scale clip-seq data. Nucleic acids research, 42(D1):D92–D97, 2014a.

Y. Li and Z. Zhang. Computational biology in microRNA. Wiley Interdisciplinary Reviews: RNA, 6(4):435–452, 2015.

Y. Li, C. Liang, K.-C. Wong, J. Luo, and Z. Zhang. Mirsynergy: detecting synergistic mirna regulatory modules by overlapping neighbourhood expansion. Bioinformatics, 30(18):2627–2635, 2014b.

P. Liu, Y. Liu, J. Luo, and Y. Li. Mirgraph: A hybrid deep learning approach to identify microrna-target interactions by integrating heterogeneous regulatory network and genomic sequences. In 2024 IEEE International Conference on Bioinformatics and Biomedicine (BIBM), pages 1028–1035. IEEE, 2024.

S. Liu, J.-H. Li, J. Wu, K.-R. Zhou, H. Zhou, J.-H. Yang, and L.-H. Qu. StarScan: a web server for scanning small RNA targets from degradome sequencing data. Nucleic Acids Research, 43:W480–W486, 2015.

S. Patel, F. Z. Peng, K. Fraser, A. D. Friedman, P. Chatterjee, and S. Yao. EvoFlow-RNA: Generating and representing non-coding RNA with a language model. Synthetic Biology, 2025.

A. Pla, X. Zhong, and S. Rayner. miRAW: A deep learning-based approach to predict microRNA targets by analyzing whole microRNA transcripts. PLOS Computational Biology, 14(7):e1006185, 2018.

M. Rehmsmeier, P. Steffen, M. Höchsmann, and R. Giegerich. Fast and effective prediction of microrna/target duplexes. Rna, 10(10):1507–1517, 2004.

S. Sammut, K. Gresova, D. Tzimotoudis, E. Marsalkova, D. Cechak, and P. Alexiou. miRBench: novel benchmark datasets for microrna binding site prediction that mitigate against prevalent microrna frequency class bias. Bioinformatics, 41(Supplement 1):i542–i551, 2025.

N. Wang, J. Bian, Y. Li, X. Li, S. Mumtaz, L. Kong, and H. Xiong. Multi-purpose RNA language modelling with motif-aware pretraining and type-guided fine-tuning. Nature Machine Intelligence, 6(5):548–557, 2024.

M. Wen, P. Cong, Z. Zhang, H. Lu, and T. Li. Deepmirtar: a deep-learning approach for predicting human mirna targets. Bioinformatics, 34(22):3781–3787, 2018.

Z. Zhang, L. Chao, R. Jin, Y. Zhang, G. Zhou, Y. Yang, Y. Yang, K. Huang, Q. Yang, and Z. Xu. Rnagenesis: foundation model for enhanced rna sequence generation and structural insights. bioRxiv, pages 2024–12, 2024.

