## Supplementary Tables and Figures for "MiRformer: a dual-transformer-encoder framework for predicting microRNA-mRNA interactions from paired sequences"

Supplementary Table S1: **Average Precision Score (APS) on miRBench test datasets.** MiRformer evaluated in zero-shot and retrained settings, compared with published methods from the miRBench benchmark ([Sammut et al., 2025], ISMB/ECCB 2025). The top section reproduces results from the miRBench paper for reference. The bottom section reports MiRformer results from this work. Best APS per column is in bold and the second best is underlined.

| **Model (Train data)** | **Klimentova2022 test** | **Hejret2023 test** | **Manakov2022 test** | **Manakov2022 left-out** |
|---|---|---|---|---|
| *Published methods (from miRBench Table 2)* | | | | |
| TargetScanCNN | 0.74 | 0.71 | 0.77 | 0.76 |
| CnnMirTarget | 0.52 | 0.53 | 0.53 | 0.51 |
| TargetNet | 0.53 | 0.58 | 0.57 | 0.58 |
| miRBind | 0.75 | 0.80 | 0.71 | 0.71 |
| miRNA_CNN_Hejret2023 | 0.74 | 0.77 | 0.71 | 0.71 |
| InteractionAwareModel | 0.66 | 0.74 | 0.69 | 0.63 |
| RNACofold | 0.67 | 0.74 | 0.63 | 0.65 |
| Random | 0.51 | 0.51 | 0.52 | 0.50 |
| *Retrained methods (from miRBench Table 3)* | | | | |
| CNN retrained (Hejret2023) | 0.77 | **0.86** | 0.79 | <u>0.78</u> |
| CNN retrained (Manakov2022) | **0.84** | <u>0.84</u> | **0.84** | **0.81** |
| *MiRformer (this work)* | | | | |
| MiRformer, TargetScan-trained (zero-shot) | 0.74 | 0.73 | 0.77 | 0.75 |
| MiRformer, Hejret2023-trained | 0.59 | 0.75 | 0.60 | 0.56 |
| MiRformer, Manakov2022-trained | <u>0.82</u> | 0.82 | <u>0.83</u> | 0.70 |

**Notes.** miRBench is an independent benchmark providing bias-corrected miRNA binding site interaction datasets derived from CLASH and chimeric eCLIP experiments. Negative pairs retain seed-complementary regions, so a model relying solely on seed presence would perform at chance level (APS $\approx$ 0.50). The Manakov2022 left-out set contains miRNA families absent from all training and other test datasets, providing a stringent test of generalization to novel miRNA families.

*Zero-shot*: MiRformer trained on TargetScan data was evaluated directly on miRBench test sets without any retraining. Its competitive APS (0.73–0.77) demonstrates that the model learned generalizable biological patterns beyond TargetScan's heuristic rules.

*Hejret2023-trained*: Training on the small Hejret2023 train set (4,084 positive pairs) led to overfitting on non-Hejret test sets, consistent with the large model capacity relative to training set size.

*Manakov2022-trained*: Training on the large Manakov2022 train set (1,253,320 positive pairs) yields APS of 0.82–0.83 on three test sets, comparable to the best CNN results in the miRBench benchmark. Lower performance on the left-out set (APS 0.70) reflects the challenge of generalizing to entirely unseen miRNA families.

### S1 Computational scalability

We profiled the inference time and peak GPU memory of MiRformer with sliding-window attention, RNAhybrid and a variant using standard full attention across mRNA lengths from 500 to 25,000 nt (Supplementary Figure S1). The full-attention variant runs out of GPU memory (OOM) at longer sequences, whereas MiRformer scales linearly, confirming the theoretical $O(wL)$ complexity of the sliding-window mechanism. The full-attention variant exhibits quadratic memory growth and exceeds available GPU memory at longer sequences (marked with ×OOM), while MiRformer's memory usage grows linearly. RNAhybrid is a tool that is designed to scale to long mRNA sequences folding with short miRNA sequences [Rehmsmeier et al., 2004]. It also achieves linear time and memory complexity, and is able to achieve a lower computational overhead than MiRformer.

This is expected since RNAhybrid is a lightweight dynamic programming algorithms on CPU, while MiRformer is a large transformer model with multiple layers of attention and feedforward networks on GPU with substantially higher per-nucleotide computational cost.

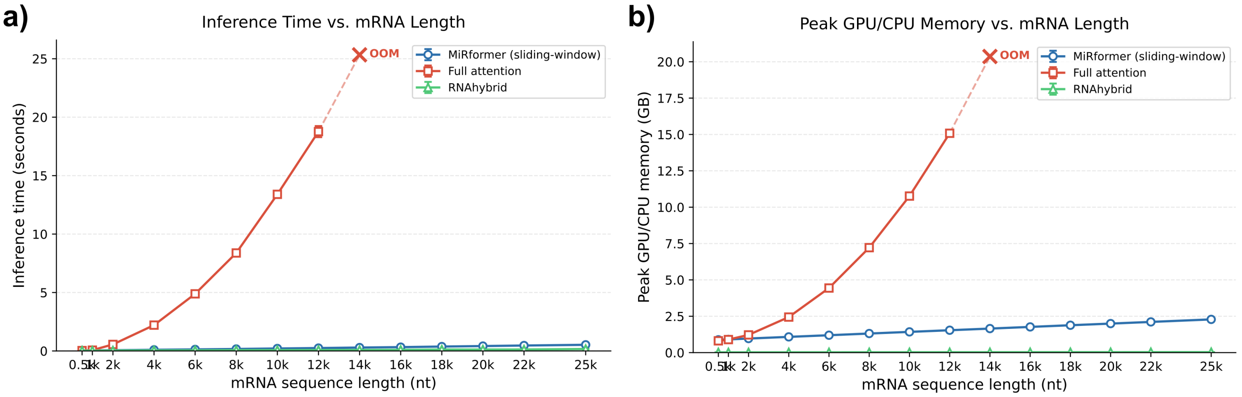


Supplementary Figure S1: a) **Inference time vs. mRNA sequence length.** Wall-clock inference time (seconds) as a function of mRNA sequence length for MiRformer with sliding-window attention (blue circles), RNAhybrid (green triangles) and a variant using standard full attention (red squares). Each point represents the median of 18 measurements (batch size = 1); error bars indicate ±1 standard deviation. b) **Peak GPU/CPU memory vs. mRNA sequence length.** Peak GPU/CPU memory usage as a function of mRNA sequence length for MiRformer with sliding-window attention (blue circles), RNAhybrid (green triangles) and a variant using standard full attention (red squares). The full-attention variant exhibits quadratic memory growth and exceeds available GPU memory at longer sequences (marked with ×OOM).

### S2 Nucleotide composition of TargetScan dataset

We examined the nucleotide composition of the 500-nt TargetScan dataset—consisting of 257,090 positive pairs and 264,986 negative pairs—by comparing G/C and A/U content and the frequencies of all 6-mers between positive and negative sets (Supplementary Figure S2). Although the distributions of G/C and A/U content differ significantly (permutation p-value $= 1.0 \times 10^{-4}$) between positives and negatives, the large overlap in their histograms (OVL = 0.81) suggests that base composition contributes only modestly to distinguishing positive from negative mRNA sequences. Moreover, the cosine similarity of approximately 0.98 between positive and negative 6-mer frequency profiles, together with the strong concentration of relative $\log_2$ fold changes around zero for positive-versus-negative 6-mer frequencies, indicates that motif composition does not act as a confounding factor (Supplementary Figure S3).

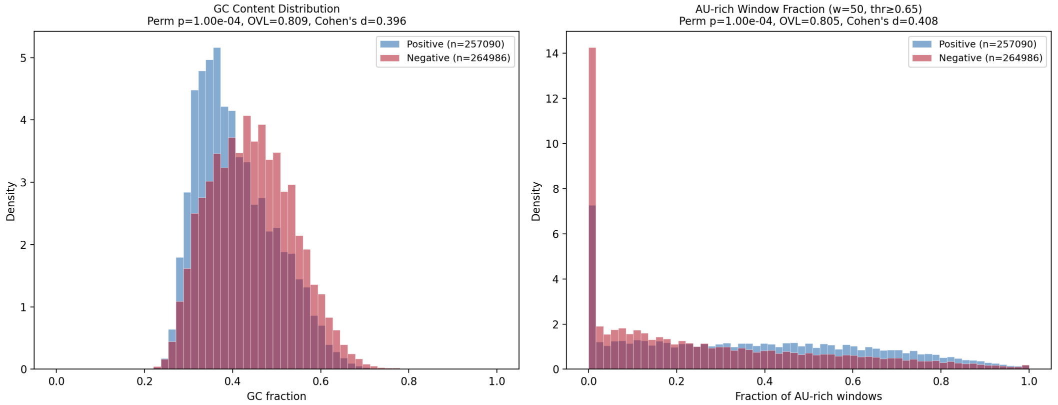


Supplementary Figure S2: **Nucleotide composition of positive and negative mRNA sequences.** Distribution of G/C content (left) and A/U content (right) in positive and negative mRNA sequences from the TargetScan 500 nt training set. Perm p indicates p-values after permutation test (10K shuffles on n=2,000 subsample). Overlap coefficient — shared area under the two histograms (1 = identical). Cohen's indicates the standardized effect size.

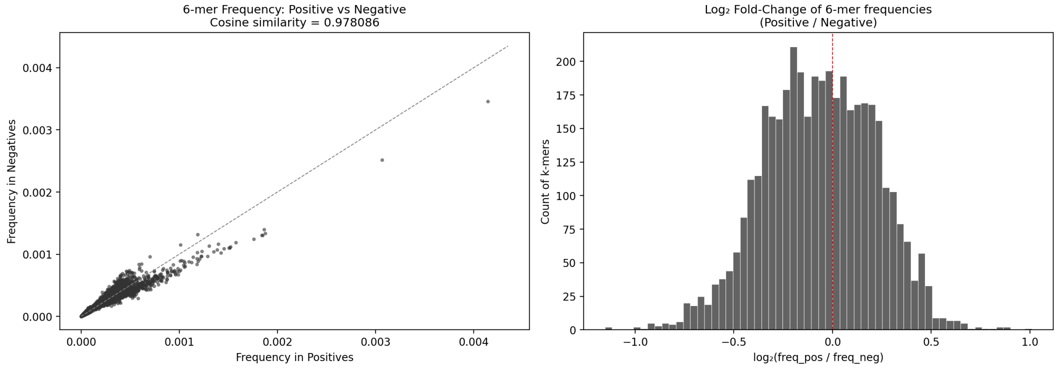


Supplementary Figure S3: **Overall 6-mer frequency spectrum in positive and negative mRNA sequences.** (Left) Scatter plot of 6-mer motif frequency in positive vs negative. Cosine similarity shows the similarity of the two distributions. (Right) Relative $\log_2$ fold change of 6-mer motif frequency in positive mRNA sequences over that in the negative. $\log_2$FC measures how much more (or less) frequent a k-mer is in one group versus the other, on a logarithmic scale:$\log_2 \mathrm{FC} = \log_2 \left( \frac{\mathrm{freq_{pos}}}{\mathrm{freq_{neg}}} \right)$.

#### S3 Investigate pairing Degradome-seq cleavage sites to TargetScan-predicted seed matches

Although the two datasets were originally unpaired, we attempted to link Degradome-seq cleavage sites to TargetScan seed matches by identifying matching pairs of miRNA name and transcript ID. This simple pairing by identifiers yielded a 54.4% match rate; however, it does not guarantee correct cleavage site coordinates within target transcripts.

To establish reliable coordinate-level correspondence, we converted genomic cleavage site coordinates to TargetScan 3'UTR transcript coordinates using the following procedure:

1. For each transcript in the Degradome-seq dataset, aggregate all associated TargetScan UTR blocks and sort them in ascending order for '+' strand transcripts and in descending order for '−' strand transcripts. This reconstruction yields the 5'-to-3' transcript structure used by TargetScan.

2. Assign each cleavage site to a TargetScan transcript block by identifying the block that contains its genomic position. If no block encompasses the site, the entry is treated as a mismatch, likely arising from isoform differences, reference version discrepancies, or annotation errors.

3. Perform a round-trip validation by converting the mapped TargetScan transcript position back to a genomic coordinate; the resulting genomic position must coincide with the original cleavage site.

4. Require a separation of only 0-5 nt between the mapped cleavage site and the nearest TargetScan seed region.

By following these stringent criteria, we identified 354 paired cleavage sites with corresponding TargetScan seed regions from the 210k Human Degradome-seq samples, corresponding to a match rate of 0.17% (Supplementary FigureS4). The low match rate is primarily attributable to two factors:

**1. Unpaired miRNA–mRNA combinations (dominant cause).** Since TargetScan annotations are restricted to 3'UTR regions, we first filtered Degradome-seq to 3'UTR-mapping sites, retaining only 63,332 of 209,668 rows (30.2%). Among these 3'UTR rows, 55.1% involve miRNA–mRNA pairs were absent from TargetScan. This is the largest source of mismatch and reflects the fundamentally independent nature of the two datasets: TargetScan predicts interactions computationally based on seed complementarity and conservation, while Degradome-seq captures experimentally observed cleavage events—including those mediated by non-canonical binding that TargetScan does not predict, and those in tissues or conditions not represented in TargetScan's conservation analysis. An additional 29.4% lack TargetScan UTR annotation blocks (likely due to isoform differences or reference version discrepancies), 13.0% have no predicted seed sites, and 1.4% fail the round-trip coordinate validation. Only 1.0% of 3'UTR rows (637 pairs) pass all matching criteria at any distance.

**2. Stringent distance threshold.** Among the 637 matched pairs, we required the cleavage site to fall within 5 nt of the nearest TargetScan seed region, consistent with AGO2-mediated cleavage occurring opposite miRNA positions 10–14. This biologically motivated threshold reduced the final matched set to 354 pairs (0.56% of 3'UTR rows, 0.17% of all Degradome-seq rows). Supplementary Figure S4c shows that relaxing this threshold would increase the match rate, but at the cost of including biologically less plausible seed–cleavage pairings.

These results confirm that the low match rate is primarily biological—the two datasets independently capture different aspects of miRNA regulation—rather than algorithmic. The scarcity of

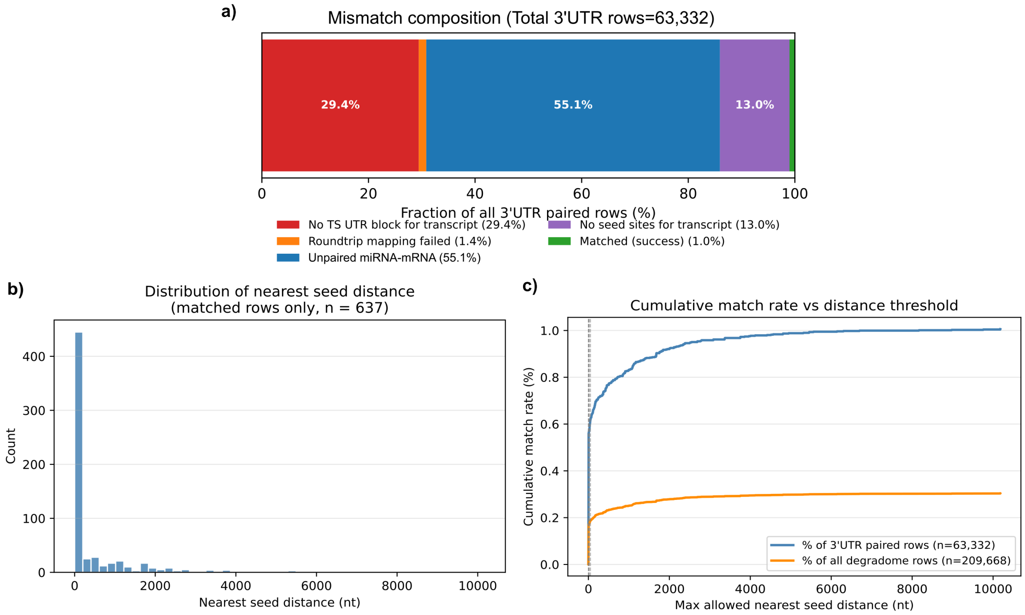


Supplementary Figure S4: **Analysis of the mismatch between Degradome-seq cleavage sites and TargetScan seed regions.** (a) Composition of matching outcomes for the 63,332 Degradome-seq cleavage sites mapping to 3'UTR regions. No TS UTR block for transcript - No TargetScan 3'UTR blocks containing the target cleavage sites; Roundtrip mapping - transcript cleavage site must be mapped back to genomic location; Unpaired miRNA-mRNA - miRNA–mRNA pairs observed in Degradome-seq that do not overlap with those predicted by TargetScan; No seed sites for transcript - no seed sites predicted by TargetScan for the mRNA; Matched (success) - miRNA-mRNA pairs that passed all constraints. (b) Distribution of nearest seed distance for the all matched pairs (n=637). (c) Cumulative match rate as a function of the maximum allowed distance threshold, shown both as a percentage of 3'UTR paired rows (blue; $n = 63{,}332$) and as a percentage of all Degradome-seq rows (orange; $n = 209{,}668$).

paired annotations motivated our approach of training MiRformer on seed recognition and cleavage prediction as separate tasks, after which the model jointly infers biologically plausible co-localized seed and cleavage sites without having been trained on paired seed–cleavage annotations.

Given the scarcity of paired annotations, we trained MiRformer on seed recognition (TargetScan) and cleavage prediction (Degradome-seq) as separate tasks, and demonstrated that the model can jointly infer biologically plausible seed and cleavage sites without training on paired seed–cleavage annotations (Fig 7).

### References

M. Rehmsmeier, P. Steffen, M. Höchsmann, and R. Giegerich. Fast and effective prediction of microrna/target duplexes. *Rna*, 10(10):1507–1517, 2004.

S. Sammut, K. Gresova, D. Tzimotoudis, E. Marsalkova, D. Cechak, and P. Alexiou. miRBench: novel benchmark datasets for microrna binding site prediction that mitigate against prevalent microrna frequency class bias. *Bioinformatics*, 41(Supplement_1):i542–i551, 2025.
